# A cellular atlas of *Astyanax mexicanus* embryogenesis reveals principles of developmental microevolution

**DOI:** 10.64898/2026.10.07.757402

**Authors:** Sergi Roig Puiggros, Goran Royer, Julien Prados, François Agnès, Julien Leclercq, Denis Jabaudon, Jorge Torres-Paz, Sylvie Rétaux

## Abstract

The evolution of development is the primary mechanism by which morphological and functional diversity is generated. Comparing the developmental trajectories of single cells in entire embryos across species can reveal how gene expression is conserved and varies, as well as the molecular mechanisms underlying how cell- and tissue-specific changes in developmental regulation generate phenotypic differences. However, this perspective has not addressed the mechanisms of small-scale developmental microevolution, i.e, in closely-related species or even within a single species. The fish *Astyanax mexicanus*, which comes in two forms adapted to distinct environments — the eyed, river-dwelling morph and the eyeless, cave-adapted morph — is a remarkable model with which to investigate the evolution of gene regulation in a single species. Here, we generated and analyzed a single-nucleus RNA-seq atlas of the development of the two *Astyanax* morphs, from gastrula to swimming larva. Despite extensive changes in gene expression observed in the eight examined stages, for both ubiquitously-expressed and tissue-specific genes, and in both embryonic and extra-embryonic tissues, the temporal progression of cavefish and surface fish cells was similar in ordinal regression-based pseudotime models using 20-100 core genes. However, transcriptome-wide comparisons of gene expression waves revealed massive heterochronic shifts. In cavefish, some tissues had a majority of genes lagging behind in their transcriptional dynamics (e.g. the optic region), while in others, a majority of genes had accelerated dynamics (e.g. the mesoderm), or else, an equivalent number of genes were found in either category (e.g. the endoderm). Furthermore, cell fate acquisition and differentiation were variably affected in different cavefish lineages, with delays of ∼6 hours in the retina, ∼2 hours in the axial mesoderm and ∼none in the extra-embryonic enveloping layer. Thus, cavefish embryos progress as a mosaic of different tissues, or modules, each following its own transcriptional tempo. Because gene expression differential regulation and time shifts were more prevalent in the early stages and gradually diminished throughout development, we propose that cavefish embryos initially undergo strong decanalisation before becoming re-canalized in line with the shared *Astyanax* developmental program.

## Introduction

The question of “How do species evolve?” lies at the heart of biology. It can be explored across multiple scales, from genomic variation to complex phenotypic traits, and from micro- to macroevolutionary timescales. Developmental evolution - encompassing subtle to profound changes in the processes that shape an organism’s tissues, forms, and functions - plays a central role in generating biological diversity and evolutionary novelties across the tree of life. Recent advances in single-cell transcriptomics have uncovered developmental trajectories in model organisms with an exquisite precision (*e.g.* [1–5]) and revealed both the conservation and divergence of these trajectories at major evolutionary transitions (*e.g.* [6–8]). Yet, small-scale phenotypic evolution, i.e., in closely-related species or even within a single species, has not been addressed. To fill this gap, here we present a comparative, micro-evo-devo single-nuclei RNA-seq atlas of embryonic development in the two eco-morphs of the emerging model species *Astyanax mexicanus*: the sighted river-dwelling fish and the blind, dark-adapted cavefish [9–11]. Our analyses uncover specific and general principles of cavefish developmental evolution, including the presence of heterochrony in developmental gene expression waves and in cell fate choices. Modularity in the timing of developmental trajectories across tissues and progressive canalization - the buffering of genetic variations [12] - emerge as components of the evolutionary toolkit in this natural mutant. Our dataset can be interrogated here https://astyanax.jablab.unige.ch/ and will serve as a valuable resource for a large community of researchers interested in developmental evolution, cavefish evolution and comparative vertebrate development.

## Results & Discussion

### An atlas of *Astyanax mexicanus* development

Our goal was to compare global developmental trajectories and cell states in *Astyanax mexicanus* embryos from surface fish (SF, from Texas) and cavefish (CF, originating from the Pachón cave [13]). We profiled 92.936 cell nuclei isolated from 780 embryos, spanning eight developmental stages from 6 to 36 hours post-fertilization (hpf). This time window covers key embryonic events, from the shield stage and the onset of gastrulation to the swimming larva stage and the onset of eye degeneration, a hallmark of cavefish development (**Fig.1A**) [14]. The sampling included ∼4.000-10.000 cell nuclei/morph/stage, a sequencing effort equivalent to initial zebrafish atlases [1, 2]. To buffer the inter-individual genetic variability inherent to the natural populations of *A. mexicanus* and to focus on inter-morph differences, we collected and pooled embryos from different spawns for each stage and morph (Methods). We retrieved rare types such as *nanos3*+ germline cells (6-8 hpf), *nkx2.1la+* neuro-mesodermal progenitors or early *oca2+* melanocytes (18-36 hpf), indicating a fair representativeness of the dataset in terms of cell types. The manually annotated and integrated UMAP of *A. mexicanus* development (SF+CF, 6-36 hpf) was consistent with a progressive, stage-dependent acquisition of tissue and cell-type identity in the two morphs, fully supported by pseudo-gene markers highlighting tissues and trajectories (**Fig.1B-D** and **Suppl Fig.1**). On both integrated and stage-specific UMAPs, the three embryonic germ layers on one hand and the extra-embryonic tissues of the enveloping layer (EVL) and the yolk syncytial layer (YSL) on the other hand, clustered separately. As expected within a single species, we could not identify morph-specific clusters or cell types (**Fig.1C** and **Suppl Fig.1**). The early relative investment of SF and CF embryos into tissues and cell types was similar, except for the EVL and YSL, underrepresented in CF (**Fig.1D**). Together, these data suggest that our dataset is representative and can be interrogated for biological insights.

**Figure 1:**
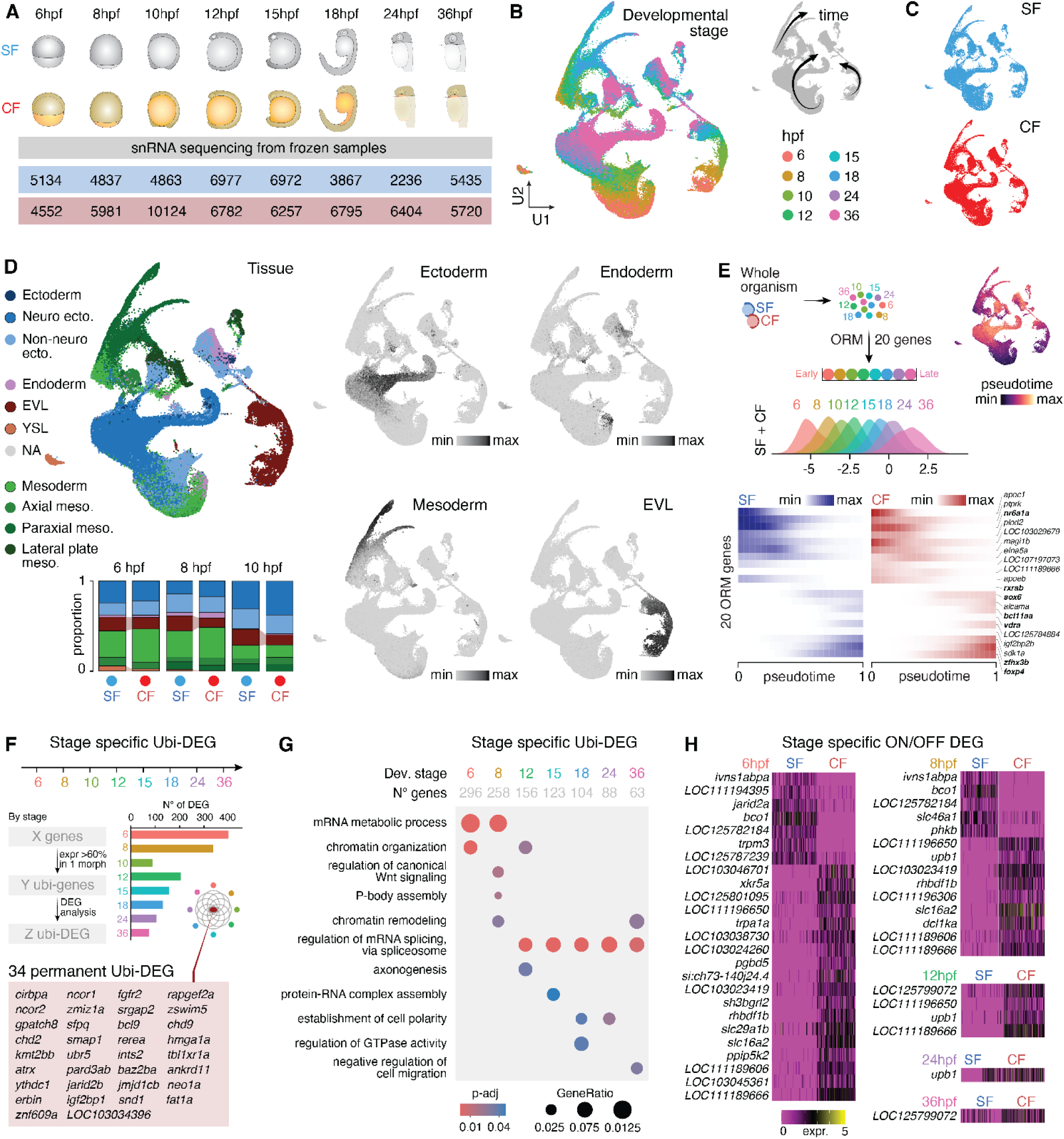
*Astyanax mexicanus* developmental cellular atlas. **A**, Stages sampled and number of nuclei sequenced in surface fish (SF, blue) and cavefish (CF, red). **B-C,** Integrated UMAPs of *Astyanax* development, color-coded by stage (B) or by morph (C). **D,** Integrated UMAPs, color-coded for tissue annotation and with pseudo-gene markers (see Methods) for the three germ layers and the extra-embryonic EVL. The bar plot shows the proportions of nuclei contributing to different tissues in SF (blue) and CF (red) embryos. **E**, Pseudotime ordering of all nuclei (all stages, SF + CF) based on an ORM with 20 genes, whose expression dynamics are shown below in SF (blue) and CF (red). Transcription factor names are bolded. The integrated UMAP on the right is color-coded by pseudotime. **F,** Histogram showing numbers of stage-specific ubi-DEGs along development (see Methods). The 34 ubiquitously-expressed genes that are permanent DEGs at the 8 stages studied are listed. **G,** Gene Ontology analysis of ubi-DEGs at the different developmental stages. **H,** ON/OFF ubi-DEGs, listed by stage, are genes expressed in all cells of one morph (black bars) but not expressed in cells of the other morph (purple color).

### The importance of ubiquitously expressed genes

To model the temporal progression of whole *Astyanax* embryos, we used ordinal regression-based pseudotime ordering ([15], Methods), which ranks cells based on their molecular temporal progression rather than embryonic age. Between 6 hpf and 36 hpf, all tissues (SF + CF pooled) could be ranked using just 20 core genes, mostly corresponding to ubiquitously-expressed genes (ubi-genes) that were progressively turned-on or -off along the 8 developmental stages (**Fig.1E** and **Suppl Fig.2**). These 20 genes (7 being transcription factors) were sufficient to capture a shared developmental trajectory across the entire organism and in the two morphs (**Fig.1E**, bottom). We propose that these genes constitute the backbone of *A. mexicanus* embryogenesis, a “core program” at single-cell resolution, and on top of which morph-specific, tissue-specific and time-specific variations can be added.

To further explore the “ubiquitous”, or generic *A. mexicanus* developmental program, we asked whether some ubi-genes (defined by expression in more than 60% of the cells in at least one morph, per stage) were differentially expressed (DEG) between SF and CF. The number of ubi-DEGs was higher at early stages (n=400 at 6 hpf) and progressively decreased through embryogenesis (n=76 at 36 hpf) (**Fig.1F** and **Suppl Table 1**). Gene Ontology (GO) enrichment analysis showed that ubi-DEGs mostly relate to mRNA processing, chromatin organization and signaling (**Fig.1G**). Further, among ubi-DEGs we identified ubi-ON/OFF genes, expressed in all cells of one morph but not expressed in the other, and thus corresponding to *bona fide* knockdowns in one morph (**Fig.1H** and **Suppl Table 1**). For example, the absence of beta-carotene hydroxylase *bco1* expression in CF tissues likely contributes to the accumulation of carotenoids and vitamin A and the associated yellowish coloration of CF embryos (**Fig.1A, 1H**), together with the recently described coding mutation in *bco2a* [14, 16, 17]. Conversely, several genes (n=33) were exclusively expressed in CF but not SF cells, such as the solute carrier genes *slc29a1b* and *slc16a2*, responsible for the transport of nucleosides and thyroid hormone, as well *ubp1*, a transcriptional activator involved in the regulation of transcription by RNA polymerase II (**Fig.1H**). We expect that these ubi-DEGs, including ON/OFF ubi-DEGs, are susceptible of having broad effects on the developmental homeostasis of cavefish cells, especially during early embryonic stages when they are the most numerous.

### Developmental trajectories of optic and axial mesoderm tissues

Molecular and temporal variations in optic and axial mesoderm tissue signaling, specification and morphogenesis are documented hallmarks of cavefish development [18–23]. We interrogated our atlas to discover the dynamic molecular bases of these phenotypes.

We analyzed the optic tissue between 10 hpf (eyefield in the anterior neural plate) and 36 hpf (formed eye), including the appearance of its two main derivatives, the neuro-retina and the retina pigmented epithelium (**Fig2.A-B**). In cavefish, optic nuclei represented a smaller proportion of the neurectoderm, consistent with a smaller investment in eye tissue, as expected (6.2% in CF *vs* 11.3% in SF at 18 hpf; 8.6% in CF *vs* 11.4% in SF at 24 hpf; 4% in CF *vs* 6.8% in SF at 36 hpf). We observed substantial transcriptional deregulation in optic cells, with 528 DEGs at 10hpf decreasing to 177 DEGs at 36 hpf, providing a list of novel candidates for the study of normal and defective eye development (**Fig.2C** and **Suppl Table 2**). GO term enrichment was logically related to cell-cell communication, neural morphogenesis and migration (**Fig.2C** and **Suppl Table 2**).

**Figure 2:**
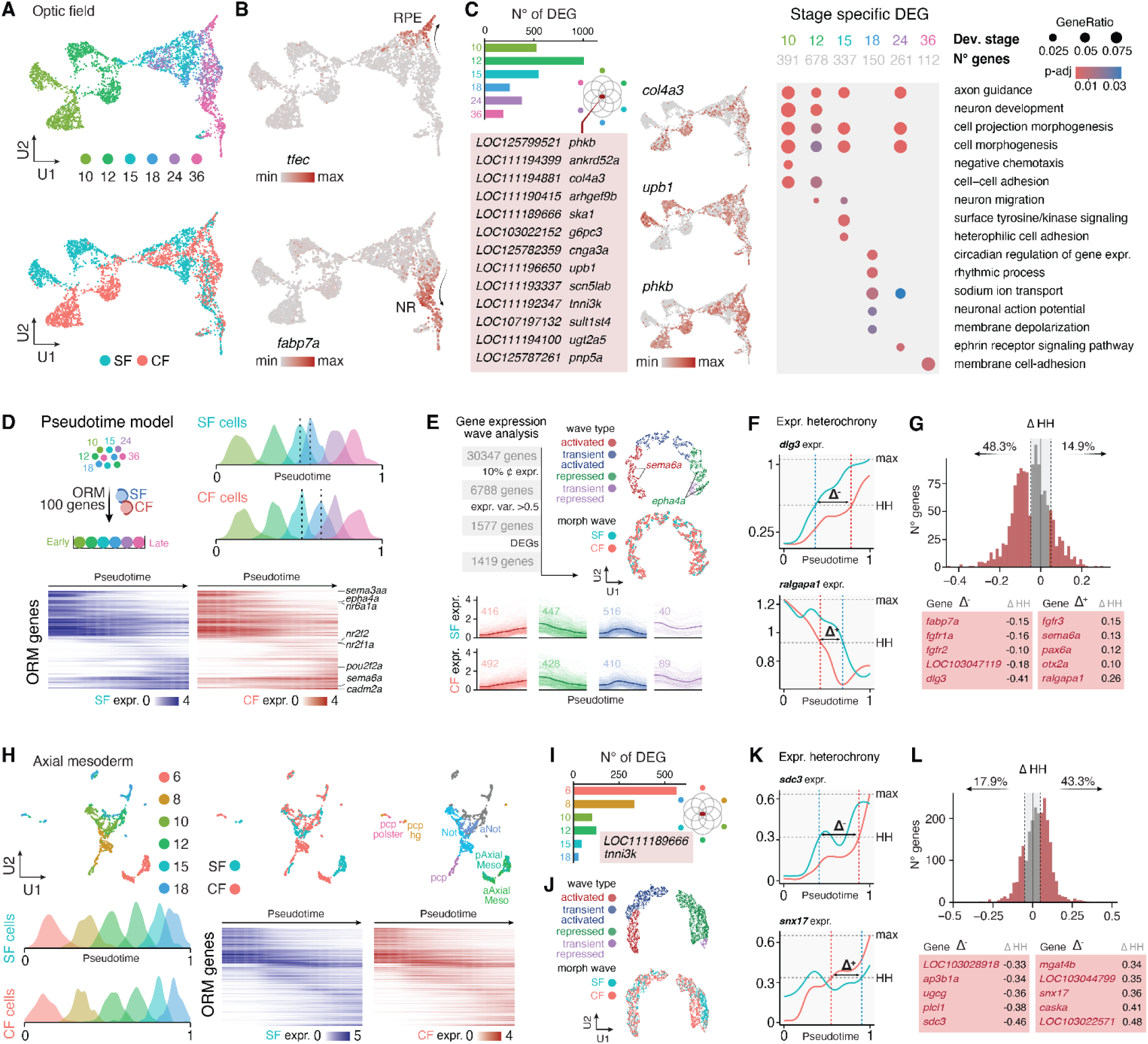
Optic and axial mesoderm tissues trajectories. **A**, UMAPs of optic tissue nuclei, color-coded by stage or by morph (SF, blue; CF, red). **B,** UMAPs showing expression of *tfec* and *fabp7*, markers of the retina pigmented epithelium (RPE) and neuroretina (NR) derivatives of the optic field, respectively. **C,** Optic tissue DEGs. The histogram shows the number of stage-specific DEGs along development (see Methods; in text and graphs, the numbers of DEGs are calculated after filtration for genes expressed in at least 10% of cells in least one morph; pvalueadj<0.05). The 26 genes that are permanent DEGs in optic nuclei at the 6 stages examined are listed, and the expression of 3 examples DEGs is plotted on the UMAP. A Gene Ontology analysis of stage-specific DEGs is shown. **D**, Pseudotime ordering and comparison of temporal progression for optic nuclei from SF and CF, based on an ORM with 100 genes, whose expression dynamics are shown in SF (blue) and CF (red). **E,** Categories of gene expression waves in the optic tissue, projected on UMAPs color-coded by wave’s categories as indicated, or by morph (see Methods). The bottom graphs show gene expression waves as a function of pseudotime for each category, in SF and CF. The number of genes in each category is indicated. **F,** The ΔHH metric as a measure of gene expression waves heterochrony. Two examples are shown. The *dlg3* expression dynamic is ‘accelerated’ in SF (blue) as compared to CF (red), hence a negative ΔHH value. Conversely, the expression dynamic of *ralgapa1* is ‘retarded’ in SF as compared to CF, hence a positive ΔHH value. **G,** Distribution of genes according to their ΔHH positive or negative value in the optic tissue, with a list of example genes. See also **Supp Fig.4. H**, UMAPs of axial mesoderm nuclei, color-coded by stage, by morph (SF, blue; CF, red), or by annotation (pcp, prechordal plate; hg, hatching gland; a/p axial meso, anterior/posterior axial mesoderm). The bottom graphs show pseudotime ordering and comparison of temporal progression for axial mesoderm cells from SF and CF, based on an ORM with 100 genes, whose expression dynamics are shown in SF (blue) and CF (red). **I,** Axial mesoderm DEGs. The histogram shows the number of stage-specific DEGs along development. See **Supp Table 3** for Gene Ontology analysis. **J,** Categories of gene expression waves in the axial mesoderm, projected on UMAPs color-coded by wave’s categories as indicated, or by morph. **F,** The ΔHH metric as a measure of gene expression waves heterochrony for two examples genes expressed in axial mesoderm. **G,** Distribution of genes according to their ΔHH positive or negative value in axial mesoderm nuclei, with a list of example genes. See also **Supp Fig.4**.

To test for possible shifts in developmental trajectories between the two morphs, we ordered SF and CF optic nuclei along ordinal regression-based pseudotime (model based on 100 genes; Methods), and we observed a conserved dynamic of optic development and progression of cell states in the two morphs - even though the 15-18 hpf transition appeared faster in SF (**Fig.2D** and **Suppl Fig.3**). To reconcile 1) apparent conservation of cell state dynamics along pseudotime with 2) global transcriptomic deregulation (DEGs) and 3) known delayed optic vesicle morphogenesis in cavefish [22, 24], we developed a pipeline to compare and classify transcriptome–wide gene expression waves and to measure gene expression heterochronies between the two morphs. Among the subset of 1419 genes that passed filters and were analyzed, 347 (24.5%) changed category between the ‘activated’, ‘repressed’, ‘transiently-activated’ or ‘transiently-repressed’ classes (**Fig.2E** and **Suppl Fig. 4**). Moreover, using a metric that represents the pseudotime shift between half-height of maximum gene expression in SF and CF (ΔHH, **Fig.2F**), we found that 896 genes showed expression heterochronies between the two morphs (**Fig.2G** and **Suppl Fig. 4**). Notably, for most of them (48%) the transcriptional dynamic was under brake in CF optic cells, while a few others (15%) showed an acceleration of their transcriptional wave. We conclude that optic development is heterochronic and transcriptionally delayed in cavefish - which aligns well with morphological observations.

We then applied the same strategy of analysis to the axial mesoderm tissue (prechordal plate and notochord) between 6 hpf (shield/organizer stage) through gastrulation and up to 18 hpf (**Fig2.H**). Of note, even though in embryos *notail*+ notochord precursors and *dkk1b*+ prechordal precursors already express a few identity markers and are already anatomically separated at 6 hpf [18] and 4 hpf (**Suppl Fig.1**), UMAP clustering did not distinguish the two cell type derivatives in the 6 hpf dataset (**Fig.2H**; see also below **Fig.5B**). Axial mesoderm nuclei of SF and CF clustered separately at early stages, and we identified 564 DEGs at 6 hpf, decreasing to 23 DEGs at 18 hpf between morphs, enriched in “signaling” GO terms (**Fig.2I**, **Suppl Table 3**). As was the case for optic cells, SF and CF axial mesoderm showed similar progression along the pseudotime model based on 100 genes (**Fig.2H**). Yet, a significant proportion of transcriptional gene waves were affected in cavefish (2750 genes were analyzed; 397, i.e., 14.4% changed category; in most cases the shift was from activated to transient activated) (**Fig2.J** and **Suppl Fig.4**). Moreover, using the ΔHH metric we found that more than thousand axial mesoderm genes were expressed heterochronically between the two types of embryos. Yet, the direction of the temporal shift was opposite to that of the optic tissue: in cavefish, heterochronic gene waves were more accelerated (43.3%) than delayed (17.9%) in their transcriptional dynamics (**Fig.2KL** and **Suppl Fig.4**). This result is consistent with experimental data on gene expression (*e.g.*, *dkk1b*) and gastrulation movements showing that dorsal convergence and the internalization and antero-posterior extension of axial mesoderm is advanced by ∼1 hour in cavefish compared to surface fish [18].

Altogether, these analyses in key, biologically relevant cell types uncover three transcriptional features of cavefish tissues developmental evolution. First, the global progression of cell states along shared pseudotime models including 100 core genes is similar in SF and CF, which may appear surprising but fits well with the two morphs developing synchronously [14], having diverged only very recently [25–27] and being the same species. Second, cavefish optic and axial mesoderm tissues undergo significant bidirectional heterochronies in the deployment and dynamics of their gene expression waves. Third, the global transcriptional dynamics follow opposite trends in different tissues – under brake in optic cells and under acceleration in axial mesoderm cells. This prompted us to test whether these original features, identified in two tissues for which significant biological knowledge is available, can be extended and generalized to other, less explored embryonic tissues.

### Embryonic tissues from all germ layers are heterochronic in cavefish

We first analyzed and compared the three embryonic germ layers. On the sets of genes analyzed in each germ layer (2197 in ectoderm, 2547 in mesoderm, 2520 in endoderm), changes in categories of gene expression waves between SF and CF were prominent in endoderm and mesoderm but less so in ectoderm (**Fig.3ABC** and **Suppl Fig.4**). Moreover, the transcriptional dynamic was differently affected in the three tissues: ectoderm and mesoderm had more advanced than retarded gene expression waves in cavefish (28.8% and 25% *versus* 10.7% and 14.8%, respectively), but in the endoderm the proportions were similar (23.5% *versus* 27.9%). Further comparing subsets of tissue types within germ layers, we found that neural and non-neural ectoderm behave similarly (**Fig3.DE** and **Suppl Fig.4**). Thus, the optic tissue emerges as an outlier in the whole ectoderm, showing a severely lagging transcriptional dynamic while the rest of the ectoderm is rather transcriptionally advanced. In the mesoderm on the other hand, both axial and non-axial mesoderm showed accelerated dynamics – although we note that this acceleration is more pronounced in the axial mesoderm (**Fig.3CF** and **Suppl Fig.4**). Finally, the vast majority of heterochronic shifts occurred at early pseudotime values for all tissue subsets (with a few at later stages in mesoderm and non-neural ectoderm) (asterisks in **Fig3**). This suggests that early steps of gastrulation and neurulation are more divergent than later steps of organogenesis and differentiation between the two morphs.

**Figure 3:**
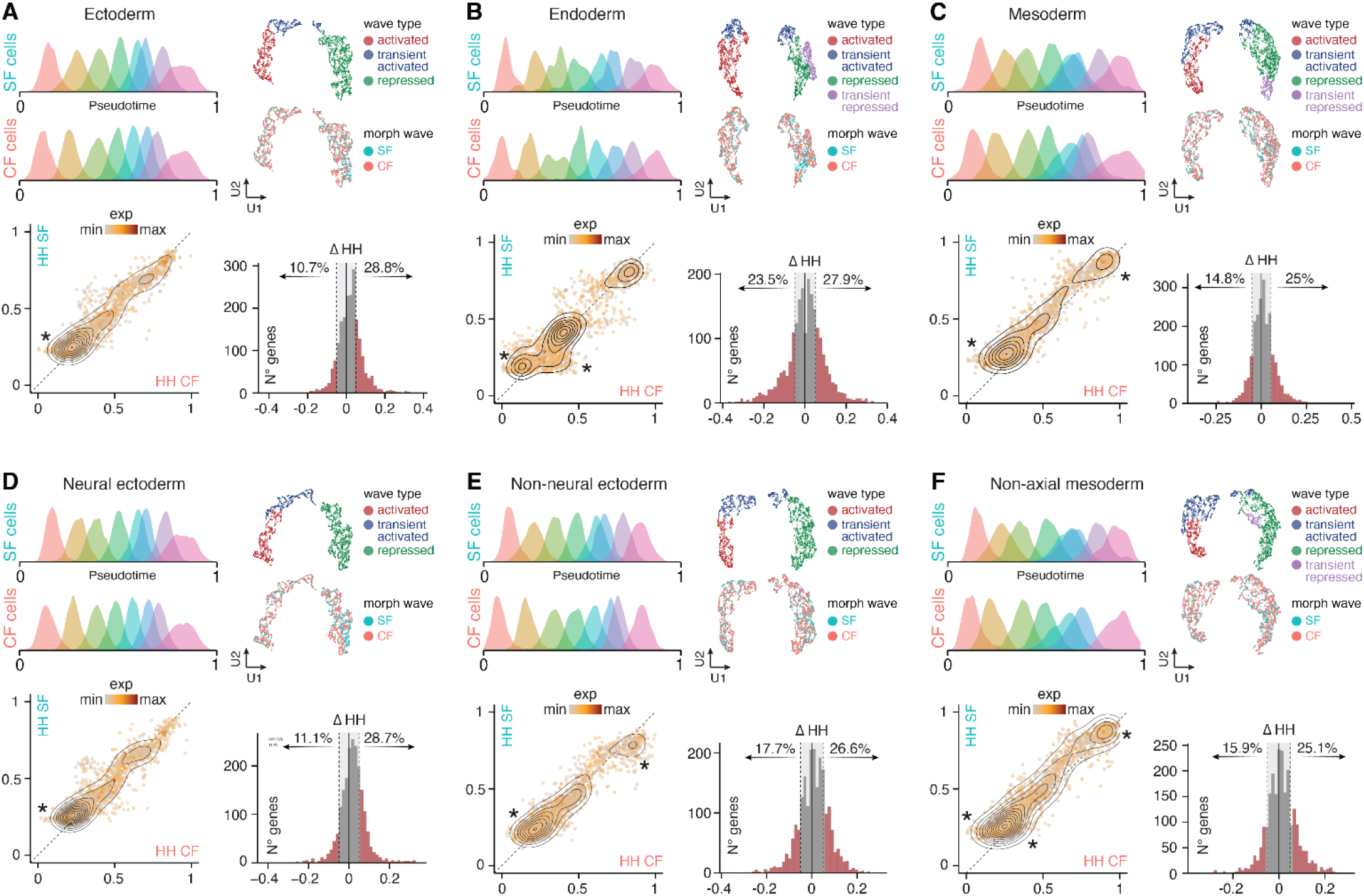
All cavefish embryonic tissues are transcriptionally heterochronic compared to surface fish. **A-F** show analyses in the indicated germ layers (ectoderm, endoderm, mesoderm) and tissue subtypes (neural ectoderm, non-neural ectoderm and non-axial mesoderm). For each tissue, the top left panel shows pseudotime ordering and comparison of temporal progression for nuclei from SF and CF, based on an ORM with 100 genes. The top right panel shows categories of gene expression waves, projected on UMAPs color-coded by wave’s categories as indicated, or by morph (see Methods). The bottom left panel show correlation plots between pseudotime values for which HH (Half Height) of maximal gene expression is reached in CF and SF. Each dot is a gene. Genes above the diagonal have accelerated dynamics in CF, whereas genes below the diagonal have delayed dynamics. The asterisks point to genes showing expression waves heterochronies mostly at early pseudotime, with some case where heterochronies occur at late pseudotimes. The bottom left panel shows the distribution of genes according to their ΔHH positive or negative value. See also **Supp Fig.4**.

Together, these findings highlight that cavefish cells follow transcriptional tempos that differ from those of surface fish, in a mosaic, tissue-specific manner. The strongly time-advanced axial mesoderm may influence through signaling and tissue interactions the pace of neural ectoderm trajectory - but not the endoderm. On the other hand, the strongly time-delayed optic tissue does not detectably disturb the rest of the future brain and behaves as an independent module (see also below). Finally, it is interesting to note that despite being considered as a mutant and ‘degenerated’ creature, cavefish embryos at early stages show globally accelerated transcriptional dynamics, which may be viewed as a gain of function.

### The extra-embryonic EVL has also evolved in cavefish

We next focused on the extra-embryonic enveloping layer, a squamous epithelial monolayer that derives from early external blastomeres. The EVL spreads over the yolk cell together with deep embryonic cells during epiboly and contributes to gastrulation, and then forms the external layer of the epidermis or periderm (**Fig.4A**, inset) [28, 29]. On the EVL UMAP, at all stages EVL nuclei from SF and CF embryos segregated in distinct and distant clusters and formed mirror images, highlighting markedly different transcriptional states despite being a single cell type sharing expression of typical and specific EVL markers such as *krt5* and *krt8* (**Fig.4AB**). We found that the derivatives of 6 hpf EVL cells are the periderm, the casquette (the *Astyanax*-specific adhesive gland [30]) and the Küpffer’s vesicle (the teleost left-right organizer; markers *sox17*, *sox32*, *dand5*) (see also **Fig.5C**). DEG analysis revealed massive differences between SF and CF EVL nuclei, especially at early stages (from ∼2838 DEGs at 6 hpf to 473 DEGs at 24 hpf) (**Fig.4C** and **Suppl Table 4**). As examples, *fgf13a* (not described for expression nor role in EVL development in zebrafish -ZFIN), LOC111189666 (an ubi-ON/OFF gene), *abca12* (expressed in zebrafish EVL and contributing to epidermis morphogenesis and keratinocyte morphology [31]) or *xkr5a* (a putative membrane transporter) were strongly deregulated in CF EVL throughout development or at specific time-windows, respectively. This further demonstrates the potential of our comparative dataset to discover new tissue-specific genes. GO enrichment analysis showed that deregulated biological processes progressed from “cell cycle” (at 6 hpf) to “keratinocyte differentiation” and “epidermis development” (at 24 hpf), in line with EVL developmental progression (**Fig.4D**). Moreover, EVL-specific DEGs were associated to intermediate filament organization and metabolic response to xenobiotics, pointing to a significant regulatory evolution of genes controlling cell structure and function in the EVL lineage between the two morphs (**Fig.4D** and **Suppl Table 4**). Of note, the Küpffer’s vesicle nuclei also displayed 1029 and 223 DEGs at 8 hpf and 10 hpf between SF and CF, respectively, mostly associated to intermediate filaments, wound healing and cell junction organization (**Suppl Table 4**). Some of these deregulations may underlie the variations in symmetry breaking events and heart asymmetry described between the two morphs [32].

**Figure 4:**
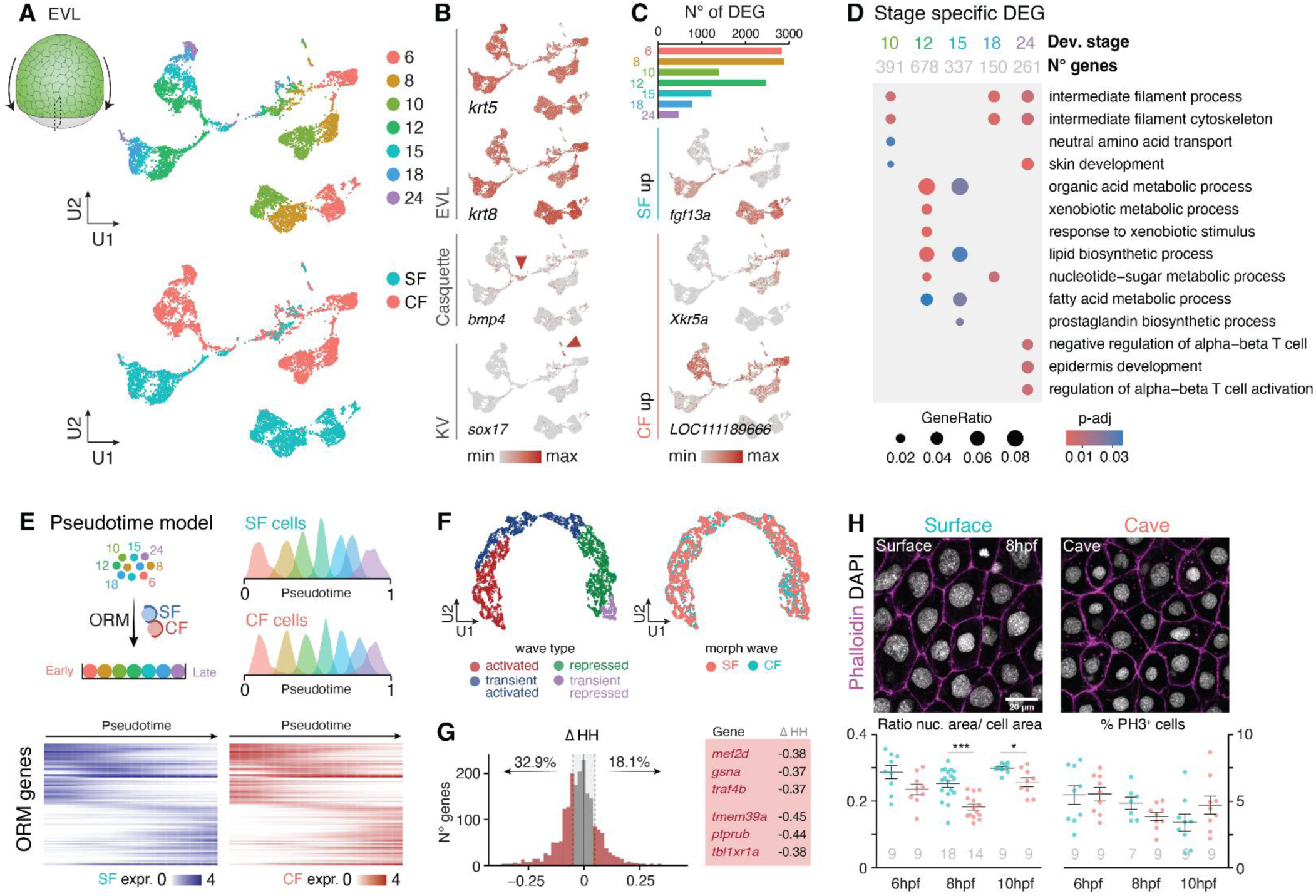
Features of the extra-embryonic enveloping layer (EVL). **A**, UMAPs of EVL nuclei, color-coded by stage or by morph (SF, blue; CF, red). The schematic shows that the EVL is the external most layer of the embryo, is composed of large cells, and spreads over the yolk together with underlying embryonic tissue during gastrulation. **B,** UMAPs showing expression of *krt5*, *krt8*, *bmp4* and *sox17*, which can be used as general or specific markers of periderm, casquette and Küpffer’s vesicle derivatives of the EVL. **C,** EVL nuclei DEGs. The histogram shows the number of stage-specific DEGs along development, and the expression of 3 example DEGs is plotted on the UMAP. **D,** Gene Ontology analysis of stage-specific DEGs. See also **Supp Table 4**. **E,** Pseudotime ordering and comparison of temporal progression for EVL nuclei from SF and CF, based on an ORM with 100 genes, whose expression dynamics are shown in SF (blue) and CF (red). **F,** Categories of gene expression waves in the EVL, projected on UMAPs color-coded by wave’s categories as indicated, or by morph. **G,** Distribution of genes according to their ΔHH positive or negative value in the EVL tissue, with a short list of examples: 3 genes whose dynamic is ‘retarded’ in CF (negative ΔHH), and 3 genes whose dynamic is ‘accelerated’ in CF (positive ΔHH). See also **Supp Fig.4. H**, Morphology of EVL cells in SF and CF embryos. Confocal images show fluorescent labelling of F-actin with phalloidin (magenta) and nuclei with DAPI (grey) at 8 hpf. The plots below show quantifications of nuclear-cytoplasmic ratio and the number of phospho-histone H3 (PH3) mitotic EVL cells at 3 different embryonic stages. * p<0.05 and *** p<0.001, Mann-Whitney tests.

Like embryonic tissues, the EVL of both morphs showed a similar progression in pseudotime according to real embryonic time, on an ordinal regression-based pseudotime model including 100 genes (**Fig.4E**). Applying our pipeline to compare gene expression waves transcriptome–wide and to quantify heterochronies, we found that EVL nuclei are also subjected to substantial variations in transcriptional dynamics (652 genes out of 2548 analyzed changed category; 25.6%) (**Fig.4F**) and display heterochronic gene expression (**Fig.4G**). However, unlike for most embryonic tissues, in CF EVL nuclei a larger proportion of heterochronic genes were lagging behind (32.9%) and fewer (18%) were accelerated in their transcriptional dynamic, as compared to SF. This finding further suggests that embryonic and extra-embryonic tissues are partly transcriptionally uncoupled through time, and developmentally mosaic.

To start exploring the anatomical correlates of the variations of EVL cells transcriptomic trajectories in CF and SF embryos we compared their morphology. We discovered significant differences in nuclear size and nucleocytoplasmic ratio (but no proliferation defect) during gastrulation, and subcellular phenotypes at larval stages (**Fig.4H** and **Suppl Fig.5**), suggesting that the variations in molecular profiles and trajectories of the cavefish EVL translate into previously unrecognized phenotypes. We propose that such divergence may at least in part result from the variations in maternal transcriptomes between SF and CF eggs [18, 28]. The EVL serves as a permeability barrier to protect the embryo from its hypotonic external environment [33], and interestingly CF have undergone major environmental shifts in physico-chemical water conditions (e.g., conductivity ∼500 µS in caves *vs* ∼1200 µS in rivers, and hypoxia) [34]. The cavefish EVL may have evolved in response to this environmental challenge.

### Transcriptional dynamics and cell fate decisions

We reasoned that the heterochronic variations in transcriptional dynamics identified above in both embryonic and extra-embryonic tissues of CF might alter the processes of cell fate decisions and differentiation. We used URD [1] to calculate transitions probabilities and construct branching trees on the ordinal pseudotime models (**Fig.5**).

**Figure 5:**
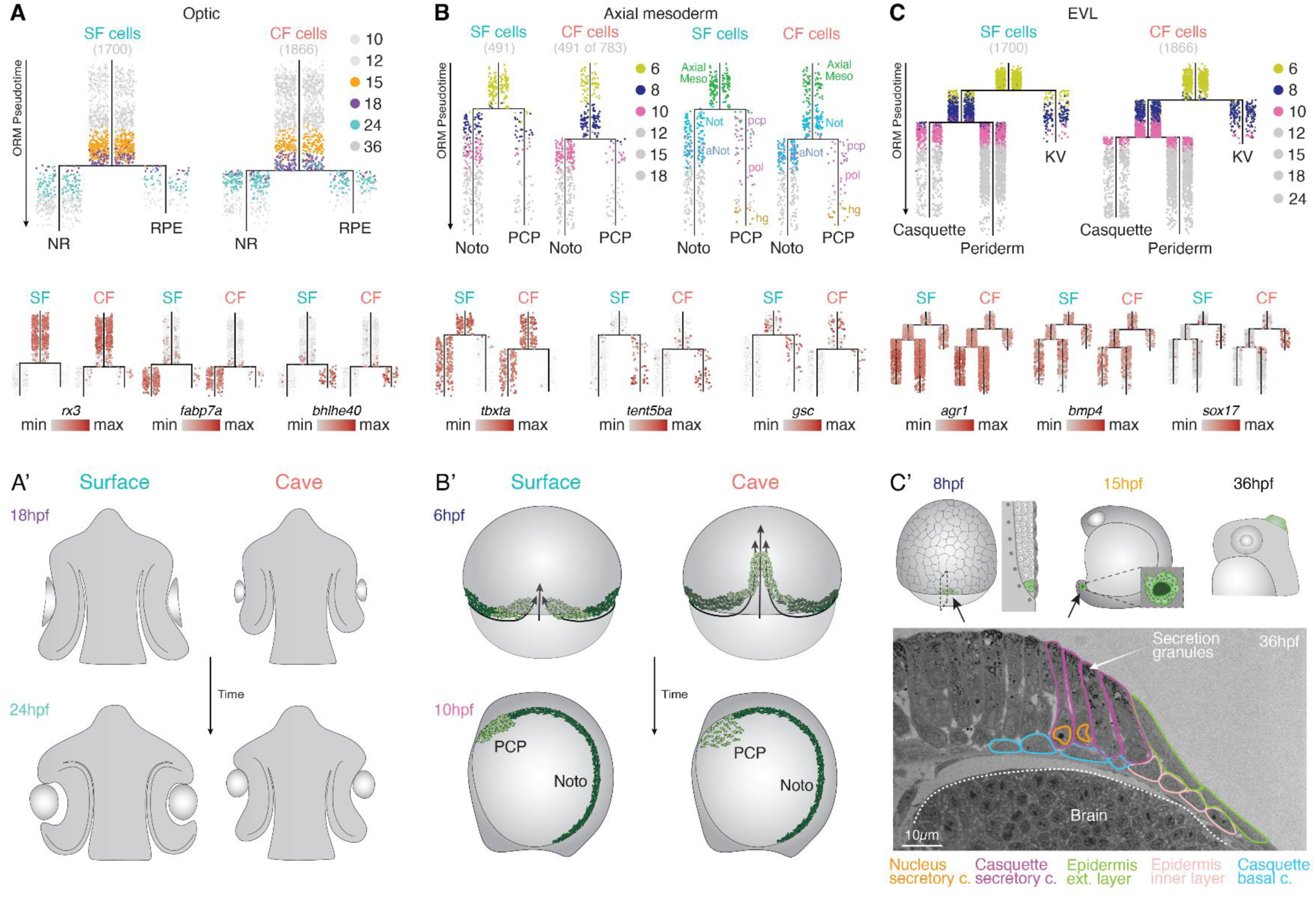
Time shift in cell fate acquisition in the optic, axial mesoderm and EVL tissues. **A**, URD tree showing the split between neuroretina (NR) and pigmented epithelium (RPE) fates for optic tissue nuclei, in SF and CF. Stages before and after branching are color-coded for better visualization. The lower panel shows cell type identity markers (*rx3*, progenitors; *fabp7*, retina; *bhlhe40*, RPE). **A’,** Schematics of relevant stages of eye morphogenesis compared in SF and CF. Adapted from [22]. The 18 hpf SF eye resembles the 24 hpf CF eye. **B,** URD tree showing the split between notochord (Noto) and prechordal plate (PCP) fates for axial mesoderm tissue nuclei, in SF and CF. On the left, stages before and after branching are color-coded for better visualization. On the right, cells are color-coded by their annotation in the atlas. Of note, the polster and the hatching gland appeared as prechordal plate derivatives on the URD trees, showing the conservation of the anterior axial mesoderm tissue lineage across fishes and confirming the accuracy of initial annotations. The lower panel shows cell type identity markers (*tbxta/notail*, notochord; *tent5a*, *gsc*, PCP). **B’**, Schematics of relevant stages of gastrulation compared in SF and CF. PCP cells (light green) and notochord cells (dark green) are shown during convergence extension (6 hpf) and after completion of migration (10 hpf). Adapted from [18]. **C,** URD tree showing the splits between periderm, casquette and Küpffer’s vesicle (KV) fates for EVL tissue nuclei, in SF and CF. Stages before and after branching are color-coded for better visualization. The lower panel shows cell type identity markers (*agr1*, *bmp4*, epidermis; *sox17*, KV). **C’,** Morphology of the different EVL derivatives in *Astyanax* embryos. At 8 hpf, the dorsal forerunner cells (DFC, green) are located at the internal dorsal margin of EVL extension. At 15 hpf, the KV (green) forms a small epithelialized vesicle in the tailbud. At 36 hpf, the casquette (green) overlies the midbrain-hindbrain boundary of the embryonic brain [30]. A semi-thin section at the level of the casquette confirms that the two layers of the casquette have distinct embryonic origins. The large, elongated secretory cells derive from the EVL and are in continuity with the external layer of the epidermis or periderm, whereas the basal cells derive from the ectoderm and are in continuity with the inner layer of the epidermis.

In the optic tissue, the split of progenitors towards neural retina or pigmented epithelium fates occurred later in CF than in SF (**Fig.5A**). Gene markers confirmed proper cell fate identities in the two branches (*fapb7* for retina, *bhlhe40*; *tfec* for pigmented epithelium). Branching occurred on cells at embryonic age mid-18 hpf in SF *versus* ∼24 hpf in CF - the 6 hours gap between the two stages sampled does not permit better resolution here. Nevertheless, in CF, the presence of 24 hpf nuclei that have not yet made their fate decision (turquoise; absent from SF tree root), together with the residual expression of the progenitor marker *rx3* in the two differentiating branches (absent from SF tree branches) suggests that cell fate choice is affected and strongly delayed in optic tissue. This aligns well with the 6 hours delay observed in cavefish embryos for optic cup invagination [22] and for pigmented epithelium morphogenesis [24], suggesting that transcriptional dynamics, cell fate decisions and morphogenetic processes are all affected and coupled in time within the optic tissue (**Fig.5A’**).

In the axial mesoderm lineage, the split towards notochord (marker *tbxta/notail*) *versus* prechordal plate differentiation (markers *gsc* and *elapor1*) occurred after 6 hpf in SF but after 8 hpf in CF, i.e., a delay of 2 hours of real embryonic time (**Fig.5B**). Interestingly, at shield stage/6 hpf, dorsal convergence and the internalization and antero-posterior extension of axial mesoderm is advanced by ∼1 hour in cavefish, and this gastrulation movement is itself uncoupled from the progression of epiboly [18]. The present data further highlight that heterochronic gene expression waves and gastrulation movements (both advanced in CF) are uncoupled in time from cell fate acquisition and differentiation processes (retarded in CF)(**Fig.5B’**). Accordingly, the axial mesoderm, which is derived from the dorsal organizer and interacts with several tissues during axis formation, can accommodate with uncoupling in the tempos of its gene expression, cell differentiation, cell movements and morphogenesis trajectories – a property that can culminate in annual killifish gastrulation [35]. In cavefish, this feature of axial mesoderm may result from both maternal influence and tweaked interactions and signaling with adjacent tissues that are themselves perturbed, and then result in modified inductive properties leading to morphological variations.

Finally, in the EVL tissue two cell fate decisions occurred. Branching towards the Küpffer’s vesicle (markers *sox17*, *sox32*), the first organ to form during vertebrate development, happened at embryonic stages late-6 hpf in SF and early-8 hpf in CF, suggesting only a ∼1 hour delay in cavefish (**Fig.5C**). Thus, at the same 6 hpf/shield stage, the time shift in fate transition is twice less in the EVL than in the axial mesoderm cells. This result also confirms from a transcriptomics trajectory standpoint that the Küpffer’s vesicle/dorsal forerunner precursor cells and the EVL derive from the same early lineage, as suggested by live imaging experiments in zebrafish [36, 37]. The second fate choice in the EVL lineage, i.e., the split between EVL/periderm and adhesive gland/casquette identity occurred at early-10 hpf in SF and mid-10 hpf in CF (**Fig.5C**), hence with a negligible embryonic time shift, consistent with initial observations that the casquette develops on the same tempo in SF and CF embryos [30]. Moreover, the EVL URD tree demonstrates that the mucus secretory cells of the *Astyanax* casquette, and thus probably those of the adhesive glands of other fish species, are yet another derivative of this extraembryonic tissue layer, in line with our histological observations (**Fig.5C’**).

In sum, even though the natural mutant CF embryos eventually develop in a proper, viable organism, their tissues undergo variations and temporal disparity in their cell fate decision process. Because time shifts in both cell fate acquisition and transcriptional waves are more or less conspicuous depending on the tissue considered, we conclude that different tissues and cell types behave as a partly temporally uncoupled mosaic, highlighting their modular nature in developing embryos. Evidence for the modularity hypothesis [38, 39] has recently emerged. Cell transplantation experiments between zebrafish and medaka embryos (3 days *versus* 9 days of embryogenesis, respectively; divergence 250 MYA) to create ‘zebrakas’ and ‘medrafish’ have highlighted temporal decoupling and autonomous developmental modules with a ‘genetic timer’, for eye organogenesis [40]. Quantitative mapping of heterochronies with single-cell genomics between these two same fish species indirectly suggests the same trend embryo-wide, as different medaka cell types show advanced, similar or delayed developmental progression in “zebrafish time” [5]. Further, genetic analyses on F2 hybrid embryos obtained from crosses between different *Oryzias* (medaka) species showed that the timing and spatial control of axis segmentation (somite formation) can be decoupled and are governed by distinct modules [41]. Our work now uncovers developmental modularity in timing as a potential source of phenotypic variation within a single species.

## Conclusions

We have discovered that both embryonic and extraembryonic tissues display major variations in their transcriptomic developmental trajectories in *Astyanax* cavefish and surface fish embryos. We identified hundreds to thousands of DEGs between the two morphs in all tissues across stages, including ubi-genes and ubi-ON/OFF genes. Despite such massive deregulations, pseudotime analyses showed that the overall progression in cell states was ‘correct’ in cavefish, with only minor shifts at some stages in all tissues. We interpret this finding by the fact that SF and CF belong to the same species and must share an *Astyanax* core program of tissue and organism development. On top of this shared program, we have identified heterochronic shifts in the deployment of gene expression waves, with most embryonic tissues (except optic) rather accelerated and the extra-embryonic EVL rather under brake in their transcriptional dynamics in cavefish. Again, such opposed dynamics do not impair the viability of embryos. Finally, we have shown shifts in the timing of cell fate decision and differentiation in different lineages, and that time shifts can be variable between tissues, from 6 hours in the extreme case of the optic cells to almost none in the case of the EVL. Therefore, cavefish embryogenesis can be seen as a mosaic of developmental modules behaving partly independently in space and time, a remarkable case-demonstration that modularity is a prerequisite for evolution through development [42]. Mosaic heterochronies may be particularly consequential in the nervous system, where brain regions and neuronal classes follow distinct developmental tempos [43, 44]. Shifts in relative timing could thus change the developmental states in which interacting partners meet, generating variations in circuit configurations and functions. Determining the extent to which such heterochronies contribute to differences in behavior and perception between surface fish and cavefish is an open line of research.

Developing embryos can adapt dynamically to genetic or environmental variation to generate organs with high precision, a process termed ‘canalization’, i.e., the tendency to buffer variation [12]. As stated by Waddington, “canalization breaks down in mutants, which may be extremely variable”, and such decanalization offers variations on which selection could act [12, 45]. Cavefish is often described as a natural mutant, so a prediction would be that its development is decanalized, which fits well with our present embryonic single-omics data. However and importantly, we note the following. First, in all tissues, the number of DEGs progressively decreases from early to late stages. Second, in all tissues, cells from SF and CF form distinct clusters at early stages and then became intermingled at later stages, between 10hpf and 15hpf, except for the EVL. Third, for all tissues, in pseudotime analyses, the 6-24 hpf nuclei could show inter-morph variations but the 36 hpf nuclei were always identical in pseudotime hence cell states in the two morphs. Fourth, heterochronic shifts in gene expression waves mostly occurred at early pseudotime stage values. Fifth, in all tissues, 36 hpf differentiated cell types were identical in the two morphs. Together these observations convergently suggest that while early cavefish embryonic tissues are highly variable and deregulated (‘decanalized’), these developmental variations are progressively and strongly buffered and ‘re-canalized’ to reach nearly indistinguishable cell types and states in the 36 hpf larvae. This hypothesis is well supported by experiments in which inhibition of the hsp90 chaperone in charge of buffering genetic variation [46] resulted in increased variance in eye size in surface fish (due to release of cryptic variations) but not in cavefish (because they are ‘re-canalized’ after the action of natural selection) [47]. Whether the tempo of canalization itself can be mosaic across cavefish tissues is an open question for future investigations.

## Data and code availability

Raw sequencing files and processed data files have been deposited at the Gene Expression Omnibus repository (BioProject ID PRJNA1540341). Download and processing instructions of processed Seurat objects from single developmental stages, tissue specific lineages can be found on the online portal associated to this manuscript (https://astyanax.jablab.unige.ch/) and on GitHub (https://github.com/goranroyer/Atlas_AST_MEX).

## Methods

### Astyanax Mexicanus embryo samples

Our colony of surface and cavefish originated from parents obtained from William Jeffery’s laboratory in 2004. The surface fish come from rivers in Texas and the cavefish come from the Pachón cave in the Mexican state of Tamaulipas. Embryos of the two morphs were obtained through *in vitro* fertilization or natural spawning after induction of breeders by altering the water temperature [48]. The embryonic development of *A. mexicanus* at 24°C is similar and synchronous for both morphs [14]. We raised and treated animals in accordance with the French and European regulations that govern the use of animals in research. SR’s authorization for the use of animals in research is 91-116. All experiments were performed on embryos at 6-36 hpf, therefore an authorization from the “Comité d’éthique en matière d’expérimentation animale” was not required.

At stages 6-8-10-12-15-18-24-36 hpf, embryos were rapidly dissected in 4°C embryo medium, with tricaine sulfate for older stages. The vitellus was removed, the head was cut posterior to the otic vesicle for stages 24-36 hpf, and samples were quickly frozen on dry ice and stored at -70°C. Depending on stages, ∼50-100 embryos or ∼30-50 heads, always originating from 2-3 independent breedings (different parents), were pooled to prepare each sample in order to increase sample’s genetic diversity and avoid the potential bias of batch effects.

### Single nuclei RNA sequencing

The collected tissue was stored at -80°C until use for single nuclei suspension generation, which we obtained through mechanical dissociation. Briefly, samples were resuspended in 1mL of ice-cold Lysis buffer (Myltenyi NUC-101) and mechanically dissociated (5-10 up & downs with a wide bore tip P1000). After 5 minutes incubation on ice, the extracted nuclei were collected by centrifugation (5 minutes at 500 x g and 4°C). This procedure was followed by two washing steps in PBS-BSA (1%), with 50U/µL of RNAse inhibitor Protector (RNAINH-RO – 3335399001). Samples were finally resuspended in 500µL of PBS-BSA (1%) and filtered through a 30µm strainer into a new low-binding 1.5mL tube. For each developmental stage, we performed simultaneous dissociations of the two morphs. 10 to 20k nuclei of each morph were loaded into the 10x Genomics Chromium Next Gem Single Cell 3’ kit v3 according to the manufacturer’s instructions. *Astyanax mexicanus* genome (AstMex3_surface - GCF_023375975.1) was used to map both morphs sequencing reads using Cellranger 6.1.1 *count*.

### Data processing and analysis

#### Quality control, integration and annotation

Low-quality cells (<1000 expressed genes) and doublets were excluded from the dataset during the first quality control. Following Seurat dedicated functions, count matrices were normalized and scaled. PCA was performed based on the top 2000 most variable genes and UMAP projections were built on the first 30 PCA dimensions and k-nearest neighbors. For the whole atlas dataset representation (**Fig.1**) and developmental stage representations (**Supp Fig.1**), we performed a CCA integration following Seurat functions. For tissue trajectory analysis, datasets were joint using the “merge” function from Seurat and processed together. For individual developmental stages, after clustering analysis, we calculated the 25-top cluster markers. Cell-type annotations were a collective effort performed based on specific marker expression combined to previous knowledge and the Zfin database (https://zfin.org/). All the annotations can be found in the online portal and the supplementary tables.

#### Pseudogene expression

Using the whole developmental atlas comprising the two morphs, we calculated the 15 top-markers using the *FindMarkers()* function of Seurat, by manually annotated germ-layer. We then computed and plotted the mean expression level for these top-markers.

#### Differentially expressed genes and GO analysis

Differentially expressed genes were calculated by stage and tissue type between morphs using the *FindMarkers()* function from Seurat. GO term enrichment was performed using *enrichGO()* function from clusterProfiler package. We ran GO term distillation using the *simplify()* function of clusterProfiler package. All the results are available in the **Supp Tables 1 to 4**.

#### Tissue-specific gene detection

Tissue specificity of gene expression was quantified using the *tau index (τ index)* [49]. For each gene, τ = Σᵢ₌₁ⁿ (1 − x^ᵢ) / (n − 1), where x^ᵢ = xᵢ / max₁≤ᵢ≤ₙ(xᵢ), xᵢ is the expression level in tissue *i*, and *n* is the number of tissues. This index ranges from 0 for ubiquitous expression to 1 for absolute specificity.

#### Pseudotime analysis

Pseudotime was built adapting ordinal regression models from the *bmrm* package as previously done [15, 50]. For the whole atlas pseudotime analysis, the analysis was based on the expression of 20 core genes. For lineage specific pseudotime calculations, we considered the expression of 100 core genes. For these, pseudotimes were calculated by tissue trajectory on both morph’s nuclei. To prevent penalization of inter-morph stage classification miss-alignment, we adjusted the model’s cost-matrix. For further details, please refer to the GitHub associated page with the full accessible code available (https://github.com/goranroyer/Atlas_AST_MEX).

#### Developmental gene wave analysis and cross-morph comparisons

As previously done [50], we modelled gene expression dynamics across the pseudotime. Briefly, for each morph and tissue lineage independently, pseudotime was truncated to a fixed range and discretized into 200 evenly spaced bins. Gene expression was smoothed using a Gaussian kernel (σ = 0.75), generating a matrix of gene expression across 200 pseudotime points representing its developmental dynamic. Before gene expression dynamic comparison between morphs, we applied three gene filtering settings: (i) genes expressed at least in 10% of the cells of one of the morphs, (ii) genes that had an expression variance (max-min) equal or higher than 0.5, (iii) genes that were not included as differentially expressed (DEGs) when applying the *FindMarkers()* Seurat function between morphs. Genes bellow these thresholds were excluded from downstream wave embedding analysis.

#### Gene wave analysis and half-height-based heterochronies

To compare gene expression dynamics across morphs, the 200-bin expression vectors were embedded using a UMAP with distance correlation. We then applied a clustering analysis on the UMAP space using Louvain community detection of a k-nearest neighbor graph. Clusters were annotated based on the average wave profile (activated, transiently activated, repressed and transiently repressed). For each gene pair (one gene, 2 morph expression profiles), we calculated the expression half-height (HH) pseudotime. It is defined as the pseudotime at which expression crosses the midpoint of its main transition. We used the surface profile as reference, after setting negative values to zero. We first identified the main expression transition by sliding a window spanning 50% of the pseudotime bins along the profile and keeping the window with the largest expression range.

Within this window, we classified the transition as ascending or descending. For monophasic profiles, this was based on the net change in expression across the window. For biphasic profiles (an internal peak), it was based on the larger variation from the window start. The HH threshold was set halfway between the peak (ascending) or trough (descending) and the opposite extreme preceding it. This threshold was applied to both morphs. For each profile, the HH pseudotime was the first bin within the window, starting from the profile’s minimum (ascending) or maximum (descending), at which expression crossed the threshold in the direction of the transition. If no crossing occurred within the window, we searched the full trajectory and kept the crossing closest to the reference crossing. A crossing was retained only if expression stayed beyond the threshold for at least 40% of the subsequent pseudotime bins. This excluded transient fluctuations. For the cave profile, HH was considered not reached if expression never spanned the threshold, or if it was already beyond the threshold at the start of the window. When the cave and surface profiles were poorly correlated (Pearson’s r < 0.75), the cave HH was taken as the sustained crossing anywhere along the trajectory that was closest to the surface HH. Genes whose profiles did not reach HH were flagged accordingly. Heterochronies were then defined as differences in HH pseudotime between morphs (HH surface – HH cave).

#### URD analysis of developmental trajectories

Developmental lineage trees were built using URD [1] bioinformatics pipeline (https://github.com/farrellja/URD). First, we computed transition probabilities between cells using *calcDM* function. The parameters *knn* and *sigma* were adjusted for each tissue analyzed. We transferred the previously calculated ORM pseudotime to morph-specific URD lineage objects. For each lineage, nuclei belonging to the earliest considered developmental stage was defined as the root, and tips were defined from the identity of the most differentiated cells. Biased random walks were performed from cells of each tip until root cells to define trajectories. Finally, URD trees were built with the *buildtree* function. Diagnostic plots were used to determine the number of pseudotime bins and the number of cells per bin to construct the tree. These parameters were adjusted for each tissue and conserved between morphs. Trees were visualized with *plotTree*, and cells labelled either by developmental stage or by gene expression level.

### Tissue processing, immunohistochemistry and quantification

Embryos were fixed in PFA 4% overnight at 4°C, washed in PBST and dechorionated. Samples were permeabilized by incubation in 0.5% Triton X-100 for 30min and then incubated in blocking solution (1% BSA, 1% DMSO, 0.5% Triton X-100 and 4% goat serum). Embryos were incubated overnight at 4°C with an anti Phospho-Histone3 primary antibody (rabbit, Sigma) diluted 1/1000 in blocking solution. Samples were washed in PBST and incubated overnight at 4°C with an anti-rabbit secondary antibody coupled with Alexa Fluor 488 diluted 1/250 (Thermo Fisher Scientific), 1 µg/mL DAPI (Sigma) and 1 µg/mL Phalloidin-iFluor 647 (Abcam). Samples were washed in PBST, and the animal pole was dissected and flat mounted in Vectashield (Vector Laboratories) for imaging. Samples were imaged with a confocal microscope (Leica TCS SP8). Mitotic cells were counted using Fiji ImageJ, and 2D segmentations of cell and nuclei contours were performed on Cellpose 2.0 [51]. Statistical analyses were performed on Graphpad prism 8.

## Acknowledgements

We thank Stéphane Père and the zootechnical service of the TEFOR unit for taking care of our *Astyanax* breeding colony, the iGE3 genomics and the FACS facility of the University of Geneva, and Hélène Hinaux and Jean-Michel Hermel for obtaining the casquette semi-thin section shown in Fig.5C.

## Author contributions

SR designed the study. JL, FA, SR and SRP performed the experiments. SRP, GR, JP, JTP and SR performed the analyses and interpreted the data. SRP built the figures with contributions from JTP and SR. JP designed and generated the code for the shiny app with contributions from SRP. SR wrote the manuscript. All authors read and approved the manuscript.

All authors disclose any financial or other interests related to the submitted work. There was no use of generative AI in the scientific writing of this manuscript.

